# Why Put-Up with Immunity when there is Resistance: An Excursion into the Population and Evolutionary Dynamics of Restriction–Modification and CRISPR-Cas

**DOI:** 10.1101/487132

**Authors:** James Gurney, Maroš Pleška, Bruce R. Levin

**Affiliations:** School of Biological Sciences, Georgia Institute of Technology, Atlanta, GA 30314; The Rockefeller University, New York, NY, 10065; Emory University, Atlanta, GA 30307

## Abstract

Bacteria can readily generate mutations that prevent bacteriophage (phage) adsorption and thus make bacteria resistant to infections with these viruses. Nevertheless, the majority of bacteria carry complex innate and/or adaptive immune systems: restriction-modification (RM) and CRISPR-Cas, respectively. Both RM and CRISPR-Cas are commonly assumed to have evolved and be maintained to protect bacteria from succumbing to infections with lytic phage. Using mathematical models and computer simulations, we explore the conditions, under which selection mediated by lytic phage will favor such complex innate and adaptive immune systems, as opposed to simple envelope resistance. The results of our analysis suggest that when populations of bacteria are confronted with lytic phage: (i) In the absence of immunity, resistance to even multiple bacteriophage species with independent receptors can evolve readily. (ii) RM immunity can benefit bacteria by preventing phage from invading established bacterial populations and particularly so when there are multiple bacteriophage species adsorbing to different receptors. (iii) Whether CRISPR-Cas immunity will prevail over envelope resistance depends critically on the length of the co-evolutionary arms race between the bacteria acquiring spacers and the phage generating CRISPR-escape mutants. We discuss the implications of these results in the context of the evolution and maintenance of RM and CRISPR-Cas and highlight fundamental questions that remain unanswered.

**Summary:** The two most widely used tools for manipulating and editing DNA restriction and Cas9 endonucleases both originate from studies of mechanisms that provide bacteria with immunity to infections with lytic bacteriophage (phage): restriction modification and CRISPR-Cas. Using mathematical and computer simulations, we explore the a priori conditions under which selection mediated by lytic phage will favor the evolution and maintenance of restriction-modification and CRISPR-Cas immunity in bacteria that, by mutation, can generate envelope resistance to these phage. The results of our analysis make predictions and raise testable-hypotheses about the genetic and ecological conditions under which these immune systems, rather than envelope resistance, will evolve and be maintained as the dominant mechanism of protecting bacteria from succumbing to infections with these viruses.

## Introduction

The two most widely used tools for manipulating and editing DNA: restriction and Cas9 endonucleases both originate from studies of mechanisms that provide bacteria with immunity to infections with lytic bacteriophage (phage): restriction modification (RM) originally identified in *Escherichia coli* (1, 2) and CRISPR-Cas found in *Streptococcus thermophilus* (3), respectively.

RM systems allow bacteria to distinguish self-DNA from non-self DNA by methylating specific DNA sequences called restriction sites. When phage-bearing DNA with unmethylated restriction sites infect bacteria carrying an RM system that recognizes these sites, the bacterial restriction endonuclease cleaves the phage DNA and aborts the infection (4). However, with a probability typically in the range of 10^−2^ - 10^−5^ (5), the infecting DNA can be erroneously methylated by the host’s DNA methyltransferase and thus escape restriction. Such “restriction escape” allows the infecting phage to bypass the immunity conferred by RM, resulting in progeny phage whose restriction sites specific to that RM system are methylated and thus protected from restriction by the same RM system (4). The phage thus modified can therefore freely replicate on bacteria carrying that RM system.

Because the restriction sites recognized by the majority of RM systems are relatively short (4-12 base pairs), the genomes of most phage are expected to carry multiple restriction sites. RM systems can thus be described as an innate immune systems, since they protect bacteria without the need for a prior exposure to the infecting phage. CRISPR-Cas, on the other hand, is commonly described as an adaptive immune system in that it requires prior exposure to the phage in order to protect bacteria from infection with that specific phage (6). Upon infection of a CRISPR-Cas carrying bacterium, presumably with a phage that is defective and incapable of replication (7), 23 to 55 base pairs-long DNA fragments from the infecting phage are incorporated as “spacers” between the short palindromic repeats located in the CRISPR-Cas region of the infected bacteria. When bacteria bearing such spacers are then infected by phage with DNA homologous to that of the acquired spacers, that DNA is recognized as foreign by an RNA-mediated mechanism and cut by the Cas endonuclease, which aborts the infection (8). Analogous to the adaptive immune system of vertebrates, CRISPR-Cas can thus retain memory of prior exposures to pathogens.

As is the case with RM, the protection provided by CRISPR-Cas is not absolute; phage can escape targeting by CRISPR-Cas via mutations in the spacer-homologous (protospacer) regions and generate CRISPR-escape mutants (9). Contrary to RM systems, which have no means to prevent infection once the phage escape restriction and become modified, bacteria with effective CRISPR-Cas systems can acquire new spacers from the escaped phage and regain immunity. A co-evolutionary arms race, in which bacteria acquire new spacers and phage respond by evolving protospacer mutations, ensues (10, 11). However, such arms races are not indefinite and can terminate when the bacteria acquire spacers to which the phage are unable to, or fail to, generate protospacer mutations (12-14) (For a more detailed review of RM and CRISP-Cas immunity, see (15) and (16) for a consideration of the evolution of different immune mechanisms).

On first consideration, it would seem that both RM and CRISPR-Cas evolved and are maintained by some form of phage-mediated selection. There is, however, a potential caveat to this hypothesis: envelope resistance. Bacteria can readily generate mutants that prevent (or significantly limit) adsorption of the phage by eliminating, modifying, or reducing the expression levels of the motifs (receptors), to which phage must bind before they can successfully enter the host cell (15). In most experimental cultures, such resistant mutants commonly emerge and become the dominant bacterial population, shortly after exposure to lytic phage (17-21). Unlike RM, which is readily defeated by host-mediated modification of escaped phage, and CRISPR-Cas, to which the phage can generate protospacer mutants, resistance is relatively permanent. Within short order, bacteria generate resistant mutants, to which the phage are incapable of evolving host-range mutations (22). To be sure, by modifying tail fibers or other structures used to adsorb to the bacteria, the phage can generate host-range mutants that overcome resistance by adsorbing to different receptors (23). How common this is and for how long such an arms race can continue is, however, currently unclear.

Why would bacteria evolve and maintain dedicated molecular mechanisms like RM and CRISPR-Cas to prevent phage infections if mutations leading to envelope resistance can be readily generated? Here, we address this question with the aid of mathematical and computer simulation models of the population and evolutionary dynamics of bacteria confronted with lytic phage. Based on our analysis, we make three predictions: (i) In the absence of immunity, even when confronted with multiple phage species recognizing different receptors and thereby requiring multiple independent mutations for envelope resistance, mutants resistant to all three phage species can evolve readily. (ii) RM protects established populations of bacteria from invasion with multiple phage species. If, however, the phage escape restriction and are modified, envelope resistant mutants are likely to ascend. (iii) If mutations leading to envelope resistance can be generated, the contribution of CRISPR-Cas mediated immunity to protecting the bacteria from phage depends on a number of factors, most important of which is the length of the arms race between bacteria acquiring new spacers and the phage generating escape mutants.

## Results

### I. Population dynamics of envelope resistance in the absence of immunity

We open this consideration with a simple base model describing the population dynamics of a single species of phage and two species of bacteria, in which there is neither RM, nor CRISPR-Cas immunity. This model is derived from that described in (24). Phage-sensitive and phage-resistant bacteria are present at densities *B*_*S*_ and *B*_*R*_ (cells per ml), respectively. A single population of phage is present at a density *P* (particles per ml). The sensitive and resistant bacteria grow at maximum rates *v*_*S*_ and *v*_*R*_ (per hour), respectively, scaled by a hyperbolic function of the concentration of the limiting resource *r* (µg per ml) (*ψ* (*r*) = *r*/(*r* + *k*)), where *k* (µg) is the concentration of the limiting resource, at which the bacteria grow at a half of their maximum rate (25). Resources are consumed at a rate jointly proportional to the net growth rate of the bacteria and a conversion efficiency parameter *e* (µg per cell) (26). At a rate *μ* (per cell per hour), sensitive cells generate resistant mutants (*B*_*S*_ ⟶ *B*_*R*_ mutation). For the sake of simplicity, we do not account for the possibility of a reverse mutation or phenotypic transitioning from resistance to sensitivity (27). The phage adsorb to sensitive bacteria with a rate constant *δ* (per cell per phage per hour) and produce *β* phage particles instantaneously (we neglect the latent periods). In our models, phage do not adsorb to envelope resistant bacteria, neither are the phage able to generate host-range mutants. At a rate *w* (per hour), a limiting resource enters the habitat of unit volume from a reservoir, where it is maintained at a concentration *c* (µg per ml). At the same rate, resources, bacteria and phage are removed from the habitat (26).

With these definitions and assumptions, the rates of change in the densities of bacteria, phage and the concentration of the resource are given by a set of coupled differential equations (28).

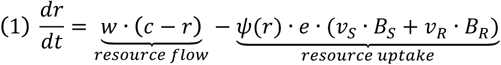

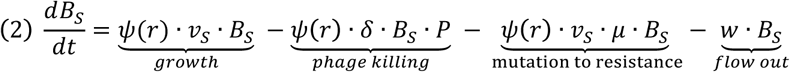

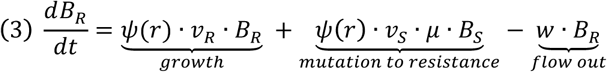

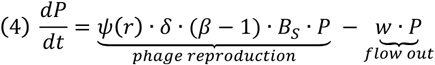

We simulated the population dynamics anticipated from this model and the models that follow using the Berkeley Madonna^™^ software. We employed parameters in the ranges estimated for *E. coli, S. thermophilus* and *Pseudomonas aeruginosa* (11, 17, 26-29) and their phage. Unlike in the above equations, where mutations are assumed to be deterministic and occur at a constant rate, mutations in our simulations were modeled stochastically. At each step of the numerical integration, a random number was drawn from a binomial distribution *D*(*n*(*t*), *μ*), where *μ* is the mutation rate and *n*(*t*) *= ψ* (*r*) *· v*_*S*_ *· B*_*S*_ *· dt* (*dt* being the step size of the numerical integration) is the number of bacterial doublings occurring at the time step *t*. We assumed a refuge density *B*_*refuge*_ = 100, below which the phage were unable to adsorb to the bacteria (30). The definitions and values of the parameters used for this model and the models to follow are listed in Table 1. The code for the Berkeley Madonna program used for this and the subsequent simulations are readily available from the authors or downloaded from www.eclf.net.

**Table 1:** Parameters and their values used in the simulations.

| Parameter | Definition | Standard values and dimensions |
| --- | --- | --- |
| $v_S, v_R, v_{RM}, v_C$ | Maximum growth rates: sensitive, resistant, RM positive, CRISPR-Cas positive, | 0.8 to 1.0 per cell per hour |
| $e$ | Conversion efficiency parameter | $5 \times 10^{-7}$ $\mu\text{g}$ per cell |
| $k$ | Monod constant | 0.25 $\mu\text{g}$ |
| $\delta$ | Phage adsorption rate | $10^{-9}$ to $10^{-7}$ per cell per phage per hour |
| $\beta$ | Burst size | 50 particles per burst |
| $\mu$ | Rate of mutation to resistance ($B_S \rightarrow B_R$) | $10^{-9}$ to $10^{-7}$ per cell per hour * |
| $c$ | Reservoir concentration of the resource | 10 to 1000 $\mu\text{g}$ per ml |
| $w$ | Flow rate | 0.2 per hour |
| $\chi$ | Phage escape (by modification) rate | $10^{-5}$ per infection * |
| $\gamma$ | Spacer acquisition rate | $10^{-7}$ to $10^{-5}$ per infection * |
| $\alpha$ | Protospacer mutation | $10^{-8}$ to $10^{-7}$ per burst * |

\* Modeled as a stochastic process

The numerical solutions of the above-presented base model are shown in Figure 1. As a result of the stochastic nature of mutations, two qualitatively different outcomes were observed: (i) Envelope resistance did not evolve. The bacteria and phage coexisted and their densities oscillated until the end of the simulation (Figure 1A). As indicated by the high concentration of the resource, the bacteria were limited by the phage. (ii) Envelope resistant mutants appeared and increased in density until fixation. The phage population declined (Figure 1B) and the bacterial population became resource-limited. We estimated the likelihood of envelope resistant bacteria evolving and becoming the dominant population of bacteria by performing repeated runs of 100 simulations, each simulating the population dynamics for 1000 hours. The results of these simulations are presented in Figure 1C. With the standard set of parameters, envelope resistant mutants dominated the bacterial population in nearly all of the runs and these populations were limited by the resource. This was also the case when resistance engendered a 20% fitness cost (*v*_*R*_ = 0.8), when the adsorption rate was low (*δ* = 10^−9^) and when the concentration of the limiting resource (and thereby the total density attainable by the bacteria) was reduced (*c* = 100 and *c* = 10). A lower rate of mutation to resistance (*μ* = 10^−9^) resulted in a reduced fraction of runs, in which resistance prevailed. Similarly, a higher rate of adsorption (*δ* = 10^−7^) resulted in only a bit more than 20% of simulations yielding bacterial populations dominated by envelope resistant bacteria. In the remaining simulations, envelope resistance did not evolve and the bacterial populations were phage-limited.

**Figure 1:**
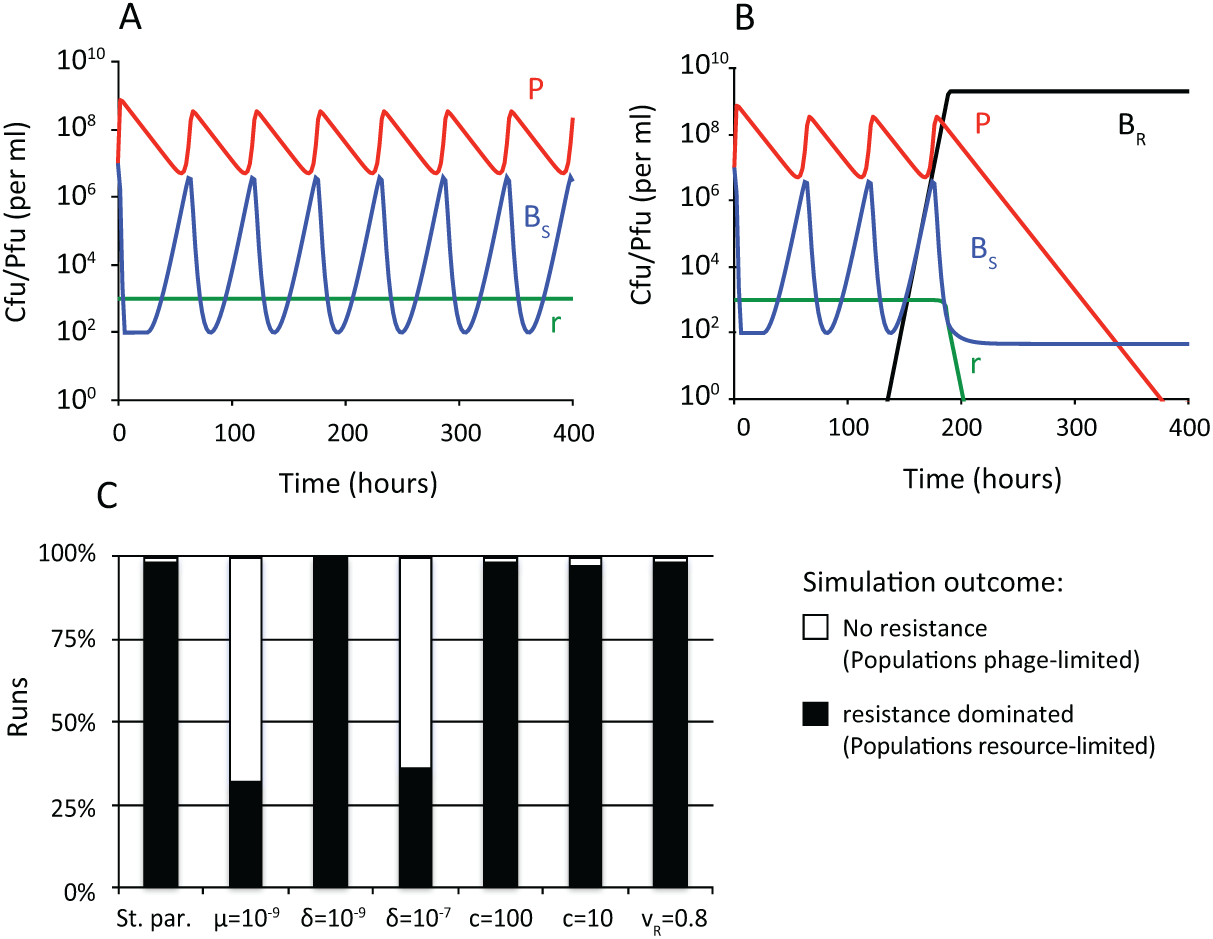
Population dynamics of bacteria and a single species of phage in the absence of immunity. Standard parameter values: *c* = 1000, *e* = 5 × 10^−7^, *k* = 0.25, *v*_*S*_= *v*_*R*_ = 1, *δ* = 10^−8^, β = 50, *μ* = 10^−8^ were used unless stated otherwise. All simulations were initiated with 10^7^ sensitive bacteria and 10^7^ phage per ml. (**A**) Changes in densities of bacteria, phage and resource concentration in a representative simulation, in which resistance did not evolve. The bacterial population is phage-limited. (**B**) Changes in densities of bacteria, phage and resource concentration in a representative simulation, in which resistance evolved. The bacterial population is resource-limited. (**C**) Outcomes at the end (t=1000) of 100 simulation runs with denoted parameter values. Populations were considered to be dominated by envelope resistant bacteria if these bacteria were the majority population. Populations were considered phage-limited if a significant concentration (> 0.9×*c*) of the resource was present and the density of the phage population exceeded that of the bacterial population.

To explore how the presence of multiple phage species affects the population dynamics of bacteria in the absence of immunity, we constructed a model analogous to that described above, but with three different phage species. The three species of phage have designations and densities *O, P*, and *Q* (particles per ml). The bacteria are of eight states with respect to resistance to these three phage: *B*_*S,S,S*_ is sensitive to all three phage; *B*_*R,S,S*_, *B*_*S,R,S*_, and *B*_*S,S,R*_ are resistant to *O, P*, and *Q*, respectively. *B*_*R,R,S*_ is resistant to *O* and *P*, but sensitive to *Q* and analogously for *B*_*R,S,R*_ and *B*_*S,R,R*_. Finally, *B*_*R,R,R*_ is resistant to all three phage. All mutations to resistance are generated at the same rate of *μ* per hour. The phage are unable to generate host range mutations to overcome resistance.

While the simulated dynamics predicted by the model with three species of phage were complex, qualitatively there were four outcomes; mutants resistant to none, one, two or all three phage species emerged and ascended to dominance before the end of the simulation. Representative dynamics of a simulation, in which mutants resistant to one phage evolved and ascended to dominance are shown in Figure 2A. In Figure 2B, we show representative dynamics of a simulation, in which resistance to all three phage (*B*_*R,R,R*_) evolved. In Figure 2C, we summarize the results of repeated runs for different sets of parameters. With the standard set of parameters, cells resistant to all three phage species evolved and dominated the bacterial population by 1000 hours in the majority of runs. In these simulations, the bacteria became limited by the resource. In the remaining 10% of the simulations, envelope resistance to only one or two of the three phage species evolved and the bacteria were limited by phage to which they remained sensitive. Comparable results were obtained when the concentration of the limiting resource was reduced (*c* = 100 and *c* = 10). When the mutation rate was low (*μ* = 10^−9^), triple-resistant bacteria ascended in only 2% of runs. Triple envelope resistance evolved and dominated all 100 simulations when the adsorption rate was low (*δ* = 10^−9^) and the opposite was true when the adsorption rate was high (*δ* = 10^−7^). Finally, triple resistance evolved in roughly a half of the simulations when resistance engendered a fitness cost (*v*_*R*_ = 0.8).

**Figure 2:**
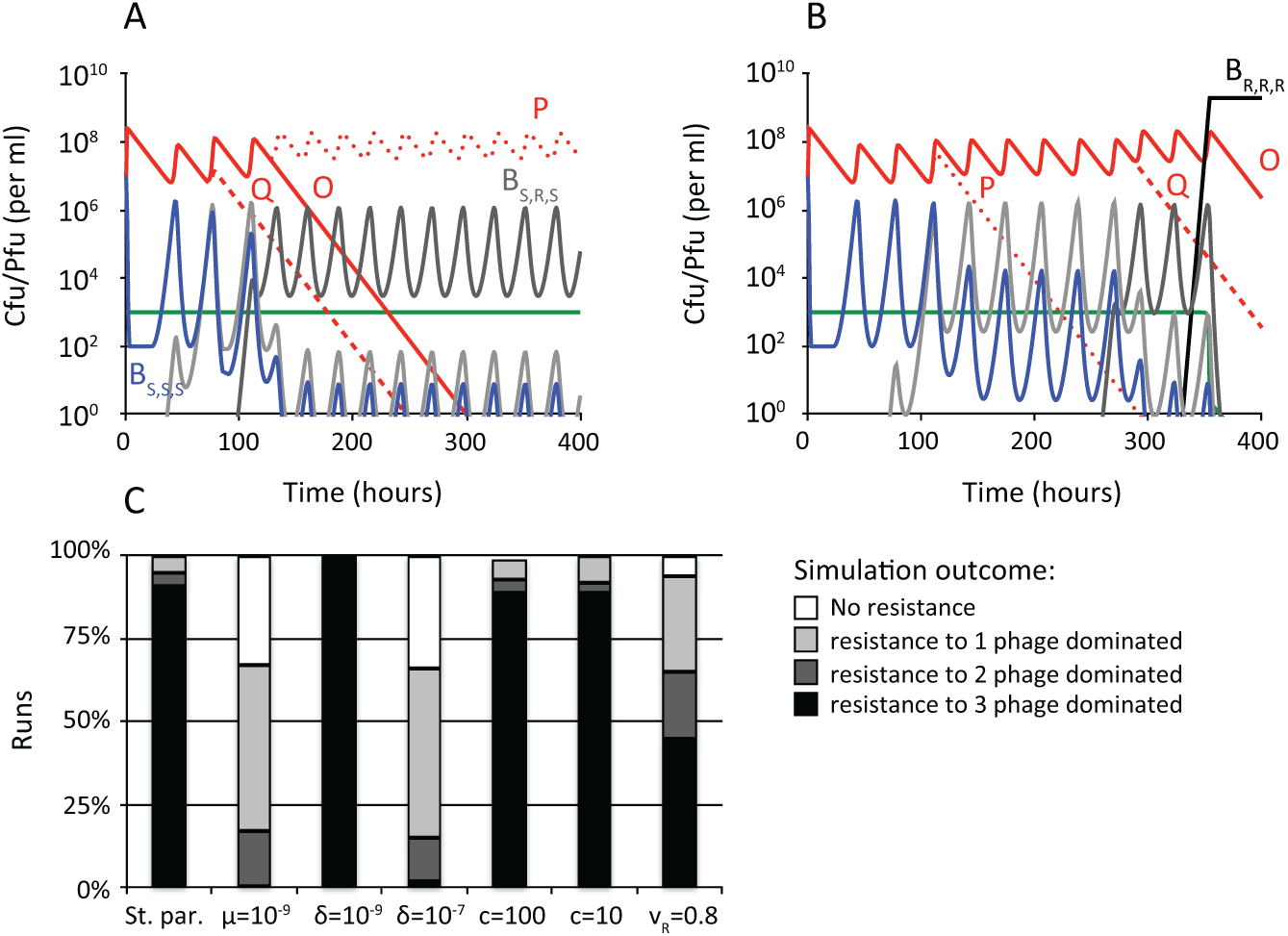
Population dynamics of bacteria and three species of phage in the absence of immunity. Standard parameter values: *c* = 1000, *e* = 5 × 10^−7^, *k* = 0.25, *v*_*S*_ = *v*_*r*_ = 1, *δ* = 10^−8^, β = 50, *μ* = 10^−8^ were used unless stated otherwise. All simulations were initiated with 10^7^ sensitive bacteria and 10^7^ phage per ml. (**A)** Changes in the densities of bacteria, phage and resource concentration in a representative simulation, in which triple resistance did not evolve. (**B)** Changes in the densities of bacteria, phage and resource concentration in a representative simulation, in which triple resistance evolved. (**C)** Outcomes at the end (t=1000) of 100 simulation runs with denoted parameter values. Populations were considered dominated by bacteria resistant to the given number of phage if these bacteria were the majority population.

### II. Population dynamics of RM immunity and envelope resistance

We constructed and analyzed a model of the population dynamics of phage and bacteria that are RM-immune by extending the above-presented model (equations 1-4) in a manner similar to what is presented in (5, 31-33). In this model, the phage can be present either in an unmodified or a modified state at densities *P* and *P*_*M*_ (particles per ml), respectively. There are two possible types of bacteria: RM-immune and resistant bacteria, which are present at densities *B*_*RM*_ and *B*_*R*_, respectively. Upon adsorption, the unmodified *P* phage are restricted by *B*_*RM*_ in the majority of cases. However, in a fraction of infections specified by the parameter *χ*, the phage escapes restriction and produces *β* modified phage particles. Modified phage can replicate on the RM immune bacteria as if these were sensitive. RM-immune bacteria can mutate into envelope resistant as described in the base model. Neither *P*, nor *P*_*M*_ phage can adsorb to resistant bacteria or generate host-range mutants. With these definitions and assumptions, the rates of change in the densities of the bacteria, phage and concentration of the resource can be described by a set of differential equations.

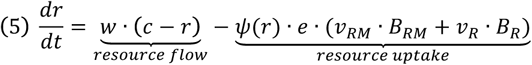

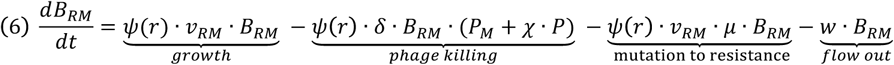

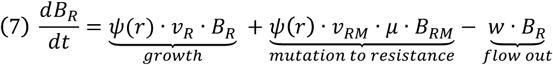

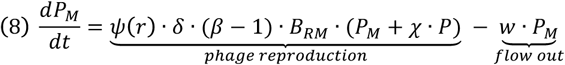

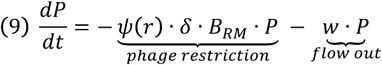

To capture the stochastic nature of mutations and phage modification, we modeled both as stochastic processes. At each time step of numerical integration, a random number corresponding to the number of escaping phage was drawn from a binomial distribution *D*(*n*(*t*), *χ*), where *χ* is the modification probability and *n*(*t*) = *ψ*(*r*) *· δ · B*_*RM*_ *· P · dt* is the number of infections occurring at the time step *t*. Mutation to resistance was modeled as described above.

In Figure 3A and Figure 3B, we present the results of two representative simulations of the population dynamics of RM-immune bacteria challenged with unmodified phage. In these simulations, we set the initial density of unmodified phage *P*(*t* = 0) to 10^5^phage particles per ml, which is equal to 1/*χ*, where *χ* = 10^−5^ is the probability of phage escaping the RM system and being modified. With this set of initial conditions, three quantitatively different results were obtained. With the standard set of parameters, the phage did not escape restriction and the modified phage were not produced in approximately one half of the simulations. In these simulations, RM-immune bacteria increased in density and the phage were eliminated by restriction (Figure 3A). In the remaining fraction of simulations, the phage did escape restriction and become modified. In these simulations, envelope resistant bacteria often emerged and fixed in the population, which was followed by phage loss (Figure 3B). In both of these cases, the populations were resource-limited. In addition to the dynamics presented in Figure 3A and Figure 3B, resistance did not ascend following the phage escape in a small number of simulations. In these simulations, the bacteria remained limited by the phage until the end of the simulation (not shown).

**Figure 3:**
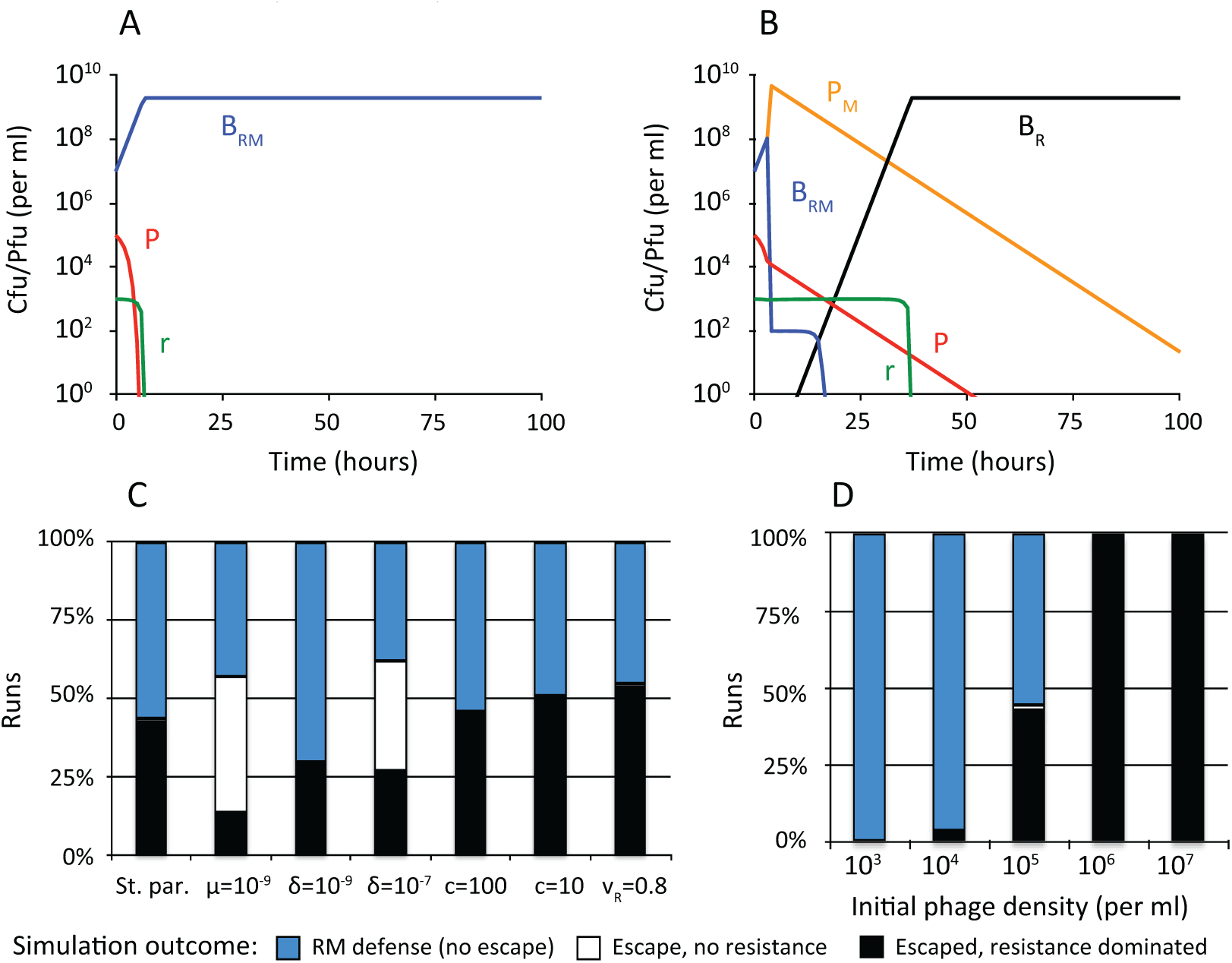
Population dynamics of RM-immune bacteria and a single species of phage. Standard parameter values: *c* = 1000, *e* = 5 × 10^−7^, *k* = 0.25, *v*_*S*_ = *v*_*R*_ = *v*_*RM*_ = 1, *δ* = 10^−8^, β = 50, *μ* = 10^−8^ were used unless stated otherwise. In all simulations, the probability of phage modification: *χ* = 10^−5^. (**A)** Changes in the densities of bacteria, phage and resource concentration in a representative simulation, in which the phage did not escape restriction and resistance did not evolve. (**B)** Changes in the densities of bacteria, phage and resource concentration in a representative simulation, in which the phage escaped restriction and resistance evolved. (**C)** Outcomes (t=1000) of 100 simulation runs with denoted parameters. (**D**) Outcomes at the end (t=1000) of 100 simulation runs with the standard parameter values and different initial densities of invading unmethylated phage. Populations were considered dominated by resistance if resistant bacteria were the majority population and analogously for RM-dominated populations. Simulations, in which the phage escaped restriction, but resistance did not evolve are shown as white bars. Simulations shown in (A), (B), and (C) were initiated with 10^7^ RM-carrying bacteria and 10^5^ unmodified phage per ml. Simulations shown in (D) were initiated with 10^7^ RM-carrying bacteria and the denoted density of the unmodified phage.

In Figure 3C we show how the key parameters of the model affect the relative likelihood of the three different outcomes. In agreement with the results shown in Figure 2C, the likelihood of resistance dominating was reduced when the mutation rate was low (*μ* = 10^−9^), or when the adsorption rate was high (*δ* = 10^−7^). Decreasing the adsorption rate (*δ* = 10^−9^) resulted in an increased likelihood of envelope resistance dominating. In all cases, the likelihood of phage escaping restriction was relatively insensitive to changing the reaction rates included in the model. The likelihood of phage escaping, on the other hand strongly depended on the initial number of phage present. The results of repeated simulations in which the initial phage density was varied are shown in Figure 3D. When the number of invading phage was low, RM immunity completely protected the bacteria from invasion and resistance never ascended. When the number of invading phage was comparable to the inverse of the modification probability *χ*, the phage invaded that RM-immune population in approximately 50% cases and this was typically followed by ascent of envelope resistance. The protective capacity of RM systems further declined as the number of invading phage increased.

We extended the above-presented model of population dynamics of RM-immune bacteria to three distinct phage species in a manner similar to the model with three species of phage and no immunity. Here, we assume that all three phage species (*O, P*, and *Q*) have an equal probability of modification *χ* = 10^−5^ and bacteria can acquire envelope resistance to these three phage species by individual mutations, each of which occurs at the same rate (*μ* = 10^−8^). The RM-immune bacteria (*B*_*RM*_) can thus, through mutation, acquire resistance to any of the three phage species and eventually become resistant to all three phage (*B*_*R,R,R*_). In these simulations, each phage species was introduced at the initial density of 10^5^ phage particles per ml.

In Figure 4, we present results of simulations with RM-immune bacteria and three, initially unmodified, phage species. Five qualitatively different outcomes were observed. In a fraction of simulations, all three phage were restricted by the bacteria carrying the RM system and envelope resistance to any of the three phage species did not evolve (Figure 4A). In the remaining fraction of simulations, one, two, or all three phage escaped restriction and became modified, which was followed by ascent of bacteria resistant to one, two or all three phage species (Figure 4B). In a small fraction of simulations, phage escape was not followed by ascent of envelope resistant mutants (representative dynamics not shown).

**Figure 4:**
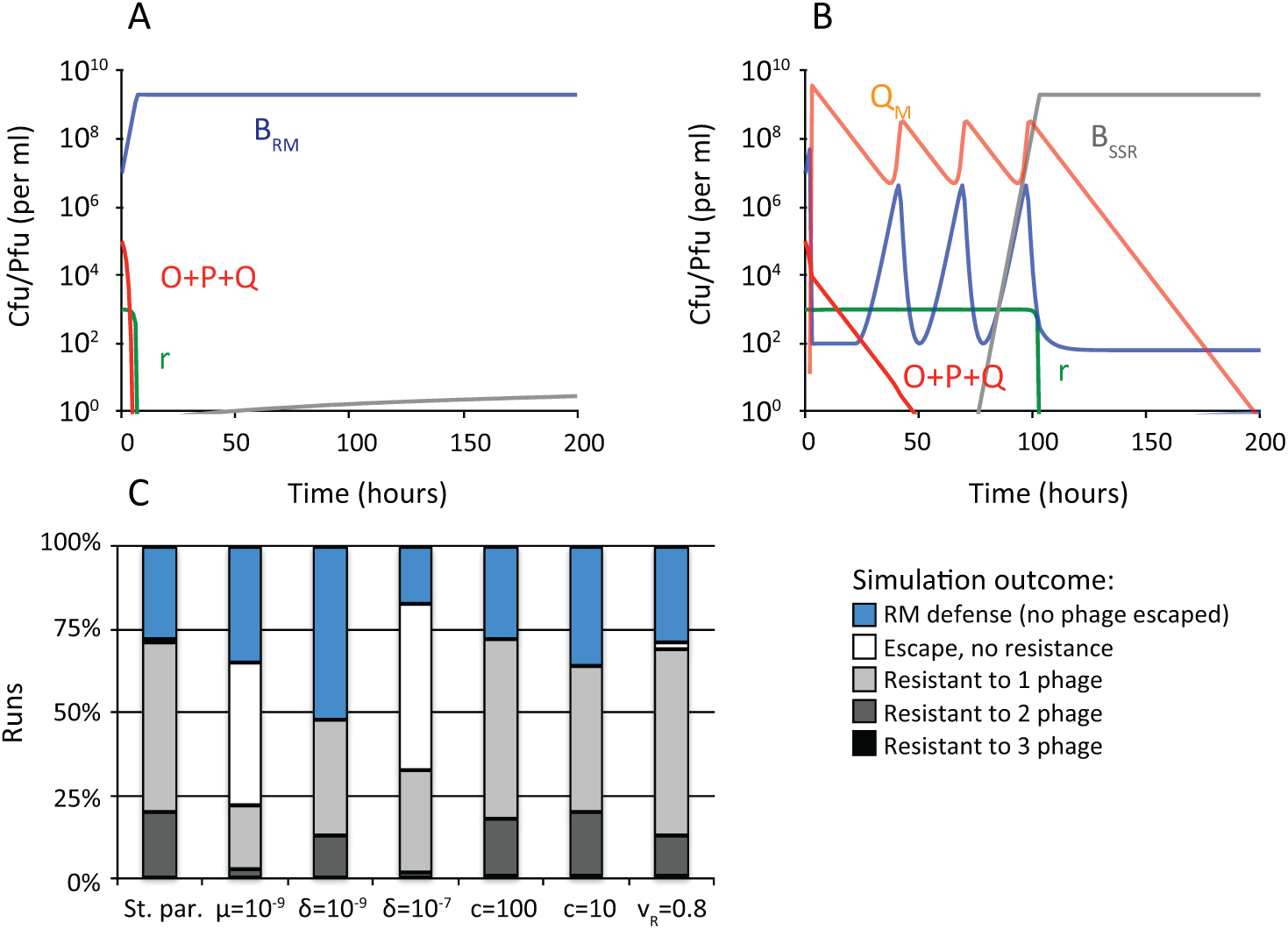
Population dynamics of RM-immune bacteria and three species of phage. Standard parameter values: *c* = 1000, *e* = 5 × 10^−7^, *k* = 0.25, *v*_*S*_ = *v*_*R*_ = *v*_*RM*_ = 1, *δ* = 10^−8^, β = 50, *μ* = 10^−8^, *χ* = 10^−5^ were used unless stated otherwise. All simulations were initiated with 10^7^ RM-immune bacteria and 10^5^ unmodified phage per ml for each phage type. (**A)** Changes in densities of bacteria, phage and resource concentration in a representative simulation, in which none of the three phage escaped restriction and resistance to any of the three phage did not evolve. (**B)** Changes in densities of bacteria, phage and resource concentration in a representative simulation, in which one phage species escaped restriction and resistance to this phage evolved. (**C)** Outcomes at the end (t=1000) of 100 simulation runs with denoted initial phage densities. Populations were considered dominated by the denoted type of bacteria if these were the majority population. Simulations, in which none of the three phage escaped restriction are shown in blue. Simulations, in which the phage escaped, but mutants resistant to even a single phage species failed to ascend are shown as white bars.

As was the case with the model of RM-immune bacteria and a single species of phage, most parameters of the model had little effect on the number of simulations, in which phage escaped restriction (Figure 4C). However, RM was more likely to prevent invasion by any of the three phage species when adsorption rate was low (*δ* = 10^−9^) and vice versa when the adsorption rate was high (*δ* = 10^−8^). Triple resistance was unlikely to ascend regardless of the parameter set used, indicating that RM represents an efficient mechanism of defending bacterial populations, in particular when multiple phage species are present.

### III. Population dynamics of CRISPR-Cas immunity and envelope resistance

The model of the population dynamics of a single phage and bacteria with CRISPR-Cas immunity described in this section is similar to the models employed in (11, 34, 35). In this model, we assume maximum seven possible types of bacteria with CRISPR-Cas immunity (*B*_0,1..6_), one type of envelope resistant bacteria (*B*_*R*_), and maximum six types of phage (*P*_0,1..5_). For bacteria with CRISPR-Cas, the integer subscripts denote the number of spacers targeting the phage. For phage, the integer subscripts denote the number of protospacer mutations carried by the phage. For example, *B*_0_ bacteria have no spacers and are susceptible to all phage. *B*_1_ bacteria have a single spacer which makes them immune to *P*_0_, but does not prevent infections by phage with one (*P*_1_) or more protospacer mutations (*P*_2..5_). In our model, *B*_6_ bacteria are immune to all phage and there is no possibility for the *P*_5_ phage to evolve an additional protospacer mutation. *B*_*R*_ is envelope resistant to all six phage and it is impossible for any phage to overcome this resistance. When CRISPR-Cas-carrying bacteria get infected with phage to which they are immune, the adsorbed phage are lost. We assume that a new spacer can be acquired by all bacteria (except for *B*_6_ and *B*_*R*_) during an infection by any phage with a probability *γ* per infection. With a probability α per successful infection, the phage acquire protospacer mutations that enable them to replicate on bacteria with the number of spacers lower than the number of protospacer mutations. All CRISPR-Cas-carrying bacteria, regardless of the number of spacers, can mutate into resistant bacteria at a rate *μ*. With these assumptions, the rates of change in the populations of bacteria, phage and the concentration of the resource are given by a set of coupled differential equations (equations 10-14).

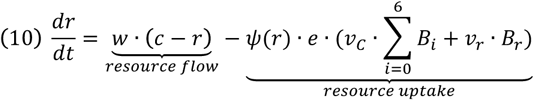

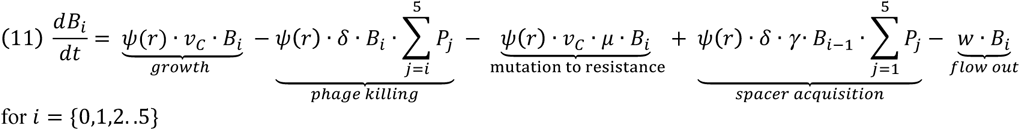

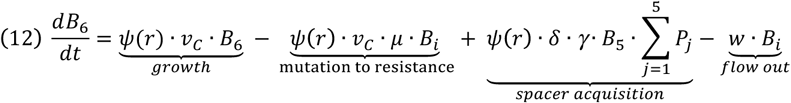

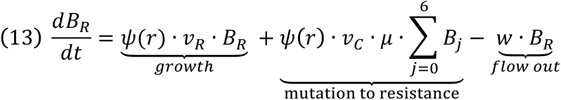

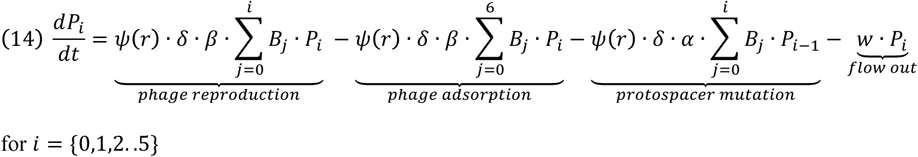

In our models, mutations from sensitivity to resistance, as well as protospacer mutations and spacer acquisition are modeled as stochastic processes. Mutations to resistance are modeled as described above in the base model with no immunity. For spacer acquisition, a random number of newly appearing *B*_*i*_ (where *i* = {1,2..6}) bacteria is drawn from a binomial distribution *D*(*n*(*t*), *γ*) at each step of the numerical integration. Here, *γ* is the spacer acquisition probability and *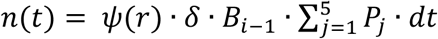* is the number of infections occurring at the time step *t*. At the same time, a random number of newly appearing *P*_*i*_ (where *i* = {1,2..5}) phage is drawn from a binomial distribution *D*(*n*(*t*), *α*), where *α* is the protospacer mutation probability and *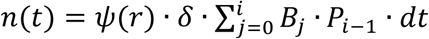* is the number of successful infections occurring at the time step *t*.

In Figure 5, we present the results of simulations obtained with the above-described model of CRISPR-Cas-immune bacteria and a single species of phage. As a result of the stochastic nature of mutation and spacer acquisition, two qualitatively different results were obtained: (i) Spacers were acquired sequentially and the phage responded with protospacer mutations until bacteria reached the state with the maximum number of spacers (*B*_6_). Envelop resistance did not ascend (Figure 5A). (ii) At some point during the coevolutionary arms race between bacteria acquiring spacers and phage evolving protospacer mutations, envelope resistant bacteria appeared and ascended to dominance. The CRISPR-Cas bacteria were maintained as a minority population (Figure 5B). In both cases, the phage were lost and the bacterial populations were limited by resources.

**Figure 5:**
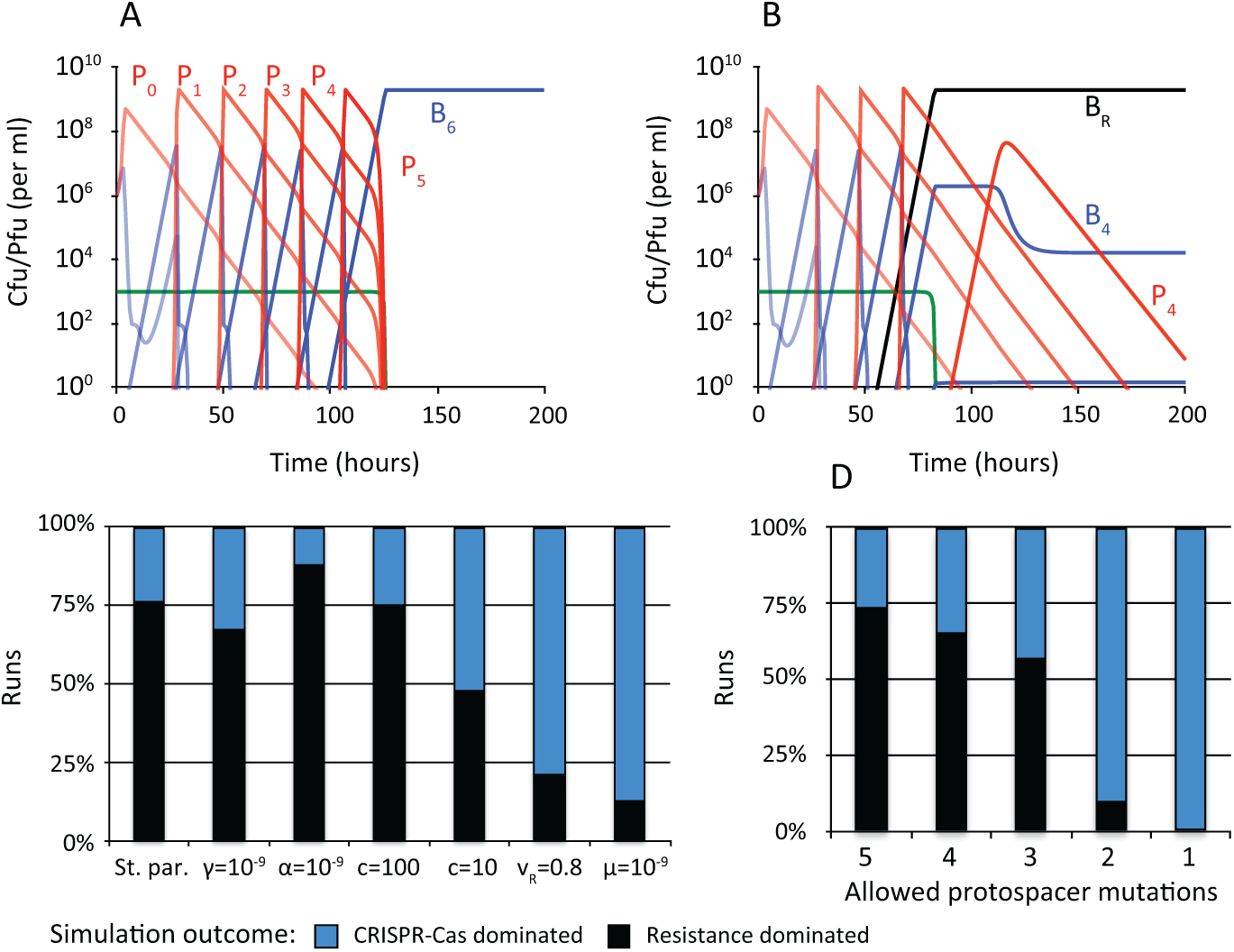
Population dynamics of bacteria with CRISPR-Cas immunity and a single species of phage. Standard parameter values: *c* = 1000, *e* = 5 × 10^−7^, *k* = 0.25, *v*_*C*_ = *v*_*R*_ = 1, *δ* = 10^−8^, β = 50, *γ* = 10^−5^, *α* = 10^−7^, *μ* = 10^−8^ were used unless stated otherwise. All simulations were initiated with 10^7^ *B*_0_ bacteria and 10^5^ *P*_0_ phage per ml. **(A**) Changes in densities of bacteria, phage and resource concentration in a representative simulation, in which resistance did not evolve and CRISPR-Cas bacteria dominated. Lines with increasing color intensity correspond to populations with increasing number of spacers (bacteria in blue) or protospacer mutations (phage in red). (**B**) Changes in densities of bacteria, phage and resource concentration in a representative simulation, in which resistance evolved and ascended to dominance. (**C**) Outcomes at the end (t=1000) of 100 simulation runs with different parameter values. In these simulations, the phage could evolve at maximum five protospacer mutations. (**D**) Outcomes at the end (t=1000) of 100 simulation runs, in which the maximum number of protospacer mutations that can be evolved by the phage was varied. The standard set of parameter values was used in these simulations.

With the standard set of parameters the bacterial populations were dominated by envelope resistant, rather than CRISPR-Cas-immune cells in 75% of simulations (Figure 5C). The relative frequency of runs in which resistance trumped (the verb) CRISPR-Cas was relatively insensitive to reducing in likelihood of acquiring a spacer (*γ* = 10^−7^), increasing the rate of protospacer mutation (*α* = 10^−5^) and reducing the concentration of the limiting resource by an order of magnitude (*c* = 100). A two-order of magnitude reduction in the resource concentration (*c* = 10), corresponding to a two-order of magnitude reduction the total cell density, led to a substantial decline in the frequency of runs, in which envelope resistance prevailed. A reduction in the number of runs in which envelope resistance dominated was also observed when resistance was associated with a significant fitness cost (*v*_*R*_ = 0.8), or when the mutation rate to resistance was low (*μ* = 10^−9^). However, it would seem that the single most important factor determining whether resistance or CRISPR-Cas immunity will prevail, is the length of the spacer-protospacer co-evolutionary arms race between bacteria and phage. To demonstrate this, we altered the maximum possible length of the co-evolutionary arms race by changing the number of maximum possible number of protospacer mutations that can be evolved by the phage before the bacteria acquire a “terminating” spacer, to which the phage cannot generate viable protospacer mutants. As the length of the spacer-protospacer co-evolutionary arms race decreased, the frequency of runs with CRISPR-Cas bacteria dominating the population increased (Figure 5D).

To explore the population dynamics of CRISPR-immunity in the presence of multiple phage species, we constructed a model capturing the population dynamics of interactions between CRISPR-Cas-immune bacteria and two distinct phage species. In this model, two rather than three phage species were considered in order to limit the number of equations. We assume that the two phage species, *P* and *Q*, have separate adsorption sites requiring two different mutations for double resistance and also genome differences great enough so that spacers acquired from phage *P* do not target phage *Q* and vice versa. The CRISPR-Cas-carrying bacteria, designated as *B*_*i,j*_, acquire spacers from these phage. The subscript *i* designates the number of spacers acquired from phage *P*, and *j* designates the number of spacers acquired from phage *Q*. The subscripts *i* and *j* can take values 0, 1, 2, 3, 4, and *R*. The subscript *R* denotes that the bacteria are envelope resistant to the respective phage (*B*_*R,j*_ bacteria are envelope resistant to *P, B*_*R,R*_ are resistant to *Q* and *B*_*R,R*_ are resistant to both *P* and *Q*). Both *P* and *Q* can acquire protospacer mutations that allow them to replicate on bacteria with the number of spacers equal to or lower than the number of protospacer mutations that these phage carry. Bacteria designated as *B*_1,0_ have one spacer for the phage *P* and none for *Q*. The phage *P*_0_ cannot replicate on these bacteria, but *P*_1_, *P*_2_, and *P*_3_, as well as all *Q* phage can. Bacteria designated as *B*_*R,*1_ are resistant to all *P* phage, irrespective of the number of protospacer mutations in their genome, but can support growth of *Q* phage with one or more protospacer mutations. Importantly, we assume that both *P* and *Q* can generate at most three protospacer mutations – i.e. they are unable to overcome immunity conferred by four spacers. The remaining assumptions are identical to those presented in the model of population dynamics with CRISPR-immunity and one phage species.

In Figure 6, we present simulation results obtained using the above-described model. In these simulations, three qualitatively different outcomes were observed. (i) Spacers were acquired sequentially for both phage and both phage responded with protospacer mutations until bacteria with CRISPR-Cas immunity against both phage emerged and ascended to dominance (Figure 6A). (ii) In the course of the spacer-protospacer coevolution, bacteria with envelope resistance against one of the phage emerged and dominated the population (representative dynamics not shown). (iii) The dominating population at the end of the simulation was envelope resistant to both phage (Figure 6B).

**Figure 6:**
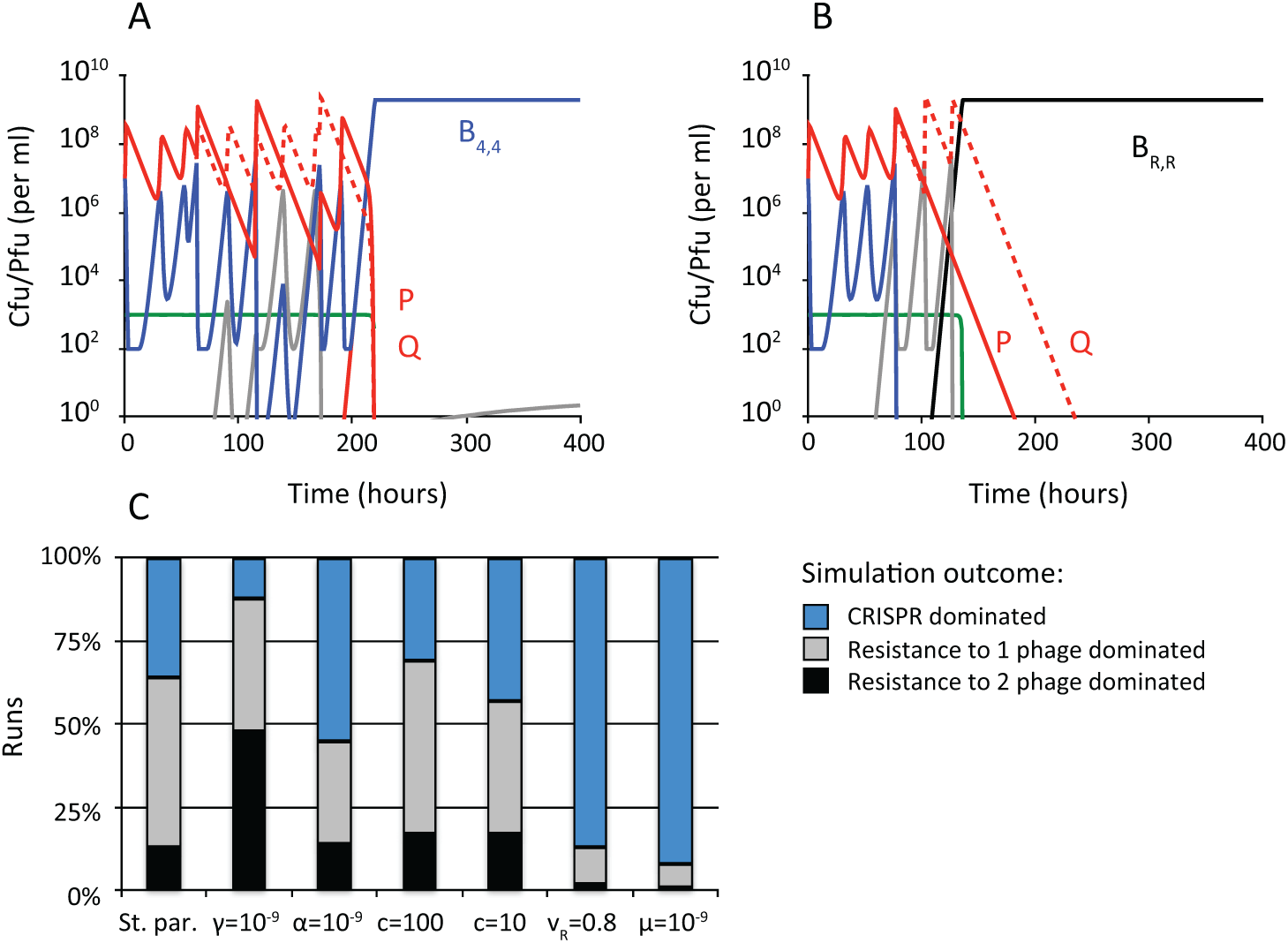
Population dynamics of bacteria with CRISPR immunity and two species of phage. Standard parameter values: *c* = 1000, *e* = 5 × 10^−7^, *k* = 0.25, *v*_*C*_ = *v*_*R*_ = 1, *δ* = 10^−8^, β = 50, *γ* = 10^−5^, *α* = 10^−7^, *μ* = 10^−8^ were used unless stated otherwise. All simulations were initiated with 10^7^ *B*_0_ bacteria 10^7^ *P*_0_ and 10^7^ *Q*_0_ phage per ml. (**A**) Changes in the densities of bacteria, phage and resource concentration in a representative simulation in which resistance did not evolve and CRISPR-immune bacteria dominated. Phage populations are shown as red lines. Bacteria with CRISPR-Cas immunity (regardless of the number of spacers) are shown in blue. Gray lines designate bacteria resistant to one phage and CRISPR-immune to the other phage. (**B**) Changes in the densities of bacteria, phage and resource concentration in a representative simulation in which resistance evolved and ascended to dominance. (**C**) Outcomes at the end (t=1000) of 100 simulations assuming denoted parameter values. Populations were considered dominated by the denoted type of bacteria if these were the majority population.

To explore how the relative likelihood of the individual outcomes depends on the parameters of the model, we performed series of repeated simulations with altered parameters (Figure 6C). With the standard set of parameters, the most frequent outcome was emergence of bacteria that were immune to one phage and resistant to the other. CRISPR-Cas immunity to both phage species prevailed more often in simulations in which resistance carried a fitness cost for the host (*v*_*R*_ = 0.8), as well as in simulations with a lowered rate of mutation to resistance (*μ* = 10^−9^). In contrast, resistance was more likely to dominate when the probability of spacer acquisition was low *γ* = 10^−7^. We observed no apparent difference in the frequency of outcomes when the resource concentration was lowered (*c* = 100 and *c* = 10). Unlike for the model with CRISPR-Cas immunity and a single species of phage, we did not explore the relationship between the length of the co-evolutionary spacer-protospacer arms race and the likelihood of resistance/CRISPR-Cas dominating. The number of equations to do so would be unwieldy. In these simulations the maximum number of protospacer mutations accessible to the phage was limited to three. It seems reasonable to assume that, similarly to what was the case for the simulations with a single species of phage, the likelihood of resistance ascending will increase with the increasing length of the co-evolutionary arms race.

## DISCUSSION

> “The most that can be expected from any model is that it can supply a useful approximation to reality: All models are wrong; some models are useful”.
>
> — Box, G. E. P.; Hunter, J. S.; Hunter, W. G. (2005), Statistics for Experimenters (2nd ed.), <u>John Wiley & Sons</u>.

We agree with this perspective by George Box and colleagues about mathematical and computer simulation models. As complex as our models may be, at best they are pale approximations of the complexity of the natural communities of bacteria and phage about which we are drawing inferences. Most importantly, our models do not consider the spatial, temporal and biological diversity of these natural communities, which, alas, is also a limitation (failing?) of most experimental systems used to study the population and evolutionary biology of bacteria and phage (for some exceptions see (36-38)) Be that as it may, these models make a number of predictions about the conditions under which RM and CRISPR-Cas immunity and envelope resistance will be favored by phage-mediated-selection; therein lies their utility.

### Envelope resistance

It is not at all surprising that our models predict that when populations of bacteria without an innate or adaptive immune system are confronted with lytic phage, there are conditions under which resistant cells will evolve and become the dominant population of bacteria. This is what has been commonly observed experimentally (17-21). Our results identify the mutation rate to resistance and the phage adsorption rate as the key parameters determining whether, in the absence of immunity, envelope resistance will evolve and the bacterial populations will become limited by resources, or on the contrary, whether resistance will not evolve and the bacterial populations will be limited by phage.

Our models further suggest that, in the absence of immunity, envelope resistance to even multiple phage species with separate adsorption sites can evolve within a relatively short time. Some 25 years ago, Korona and Levin (33) observed rapid evolution of resistance to three phage species with separate receptor sites in experimental populations of *E. coli.* The authors showed that this surprisingly fast evolution of triple resistance can be attributed to a hierarchy of phage replication, where infection with one phage suppressed the replication of the other phage (39). In the simulations presented here, albeit at a longer time scale, resistance to multiple phage species evolved even in the absence of such hierarchy. At this juncture, the question of how commonly resistance to multiple phage species will evolve in the absence of infection hierarchy remains unknow but could be addressed experimentally.

### Restriction-Modification immunity

Our models of the population dynamics of RM-immune bacteria confronted by one and three phage species make quantitative predictions about the relationship between the probability of modification, the densities of phage, and the conditions under which phage will invade and become established in populations of initially RM-immune bacteria. The qualitative prediction that RM can prevent phage from invading and becoming established in bacterial populations is intuitive. Also apparent without the aid of mathematical or computer simulation models is the recognition that unlike resistance, which is commonly specific for a single phage species, RM can offer a first line of protection from multiple phage species, with different receptor sites and thereby requiring different mutations for resistance. Our models predict that when challenged with multiple phage species, RM can reduce the effective number of phage able to invade and thus reduce the number of necessary envelope resistance mutations. Such property could be a virtue especially if we consider that unlike envelope resistance, which is commonly associated with a high fitness cost, the cost of RM is typically low (40). Importantly, even in cases in which phage do escape restriction and envelope resistance evolves, the originally RM-immune bacteria retain their RM system, envelope resistance and RM immunity are not exclusive.

Curiously, despite the ubiquity of RM systems in the bacteria and archaea and the obvious ecological importance of these innate immune systems, there have been few empirical considerations of the contribution of RM to the ecology and evolution of bacteria, archaea and the phage that prey on them. Currently, we know little about the nature and magnitude of the contribution of RM to the ecology and evolution of natural communities of bacteria, archaea and phage. From the perspective of the phage, there would be selection for evading RM, and it’s not at all clear how commonly extant phage are susceptible to RM in general. As a consequence of naturally methylated DNA, the *E. coli* phages like T2 and T4 are not susceptible to many RM systems. Moreover, many phage have other mechanisms that make them immune to RM, including a dearth of restriction sites (41-44).

### CRISPR-Cas Immunity

In the absence of resistance, CRISPR-Cas immunity will be maintained by lytic-phage-mediated selection under broad conditions. If, however, resistance can be generated by mutation, our models predict that the conditions for CRISPR-Cas to evolve and be maintained as the dominant mechanism of protecting bacteria against lytic phage are restrictive. Our models indicate that, if envelope resistance can be generated by a single mutation, the most important factor in determining whether resistance will evolve is the length of the arms race between bacteria acquiring phage-targeting spacers and phage responding by protospacer mutations. The longer this arms race, the greater the likelihood that envelope resistant mutants will ascend to dominate the bacterial population. Our models further indicate that CRISPR-Cas would be favored over envelope resistance under conditions where the mutation rate to resistance is low, or where envelope resistance carries a significant cost for the host. Such conditions could be met in environments, where presence of the receptor recognized by the lytic phage is under strong selective pressures.

In our models, we focused primarily on the question of conditions, under which complex innate and/or adaptive immunity conferred by either RM or CRISPR-Cas, respectively will prevail over simple envelope resistance. We did not explore the conditions under which the phage will be stably maintained in a population dominated by bacteria upon which the phage cannot replicate. Nor did we explore adaptive or stochastic phenotypic changes to a fluctuating environment as in (45). Whether the phage will be maintained following the ascent of resistance/immunity will depend on whether there is a mechanism maintaining a sufficiently high density of susceptible bacteria to maintain the phage. Several such mechanisms have been previously experimentally demonstrated, including a high rate of reversion from resistance (or immunity) to sensitivity, (14, 27, 46-48), a refuge of susceptible bacteria in structured habitats (30), and fitness cost of resistance (17, 22, 49-52).

It is again important to note that, despite the recent extraordinary interest in CRISPR-Cas biology, we are aware of only one empirical study that explored the conditions under which this adaptive immune system will prevail over envelope resistance when a population of bacteria with a functional CRISPR-Cas is confronted with a phage to which bacteria can generate envelope resistant mutants (53). This jointly theoretical and experimental study with *Pseudomonas aeruginosa* PA14 and a virulent mutant of the temperate, mu-like phage (DMS3-vir) presented evidence that when resistance can be generated by bacteria which also carry a functional (able to acquire spacers) CRISPR-Cas system, whether resistance or CRISPR-Cas immunity dominates depends on the nutrient composition of the media.

It would be enlightening to perform these experiments with another experimental system consisting of bacteria (or archaea) carrying functional, pick-up-spacers from phage, CRISPR-Cas system and phage, for which the bacteria can generate envelope-resistant mutants. But are there such experimental systems? To our knowledge, currently there are only two well-established, laboratory-amenable, systems, in which the bacteria with a functional CRISPR-Cas can naturally acquire spacers when infected with lytic phage: (i) *P. aeruginosa* PA14 and DMS3-vir employed in (53) and (ii) *S. thermophilus* (3, 11, 12, 14). At this time, it is not clear whether *S. thermophilus* can generate mutants with envelope resistance against to the phage at which it can readily pick up spacers.

If indeed CRISPR-Cas commonly serves as an adaptive immune system that protects bacteria from lytic phage, why are there so few experimental systems available to study this phenomenon? Could it be, that the quest of finding new experimental systems has not been intensive enough? Or could it be that despite the retrospective evidence for CRISPR-Cas functioning as an adaptive immune system – spacer sequences being homologous to phage DNA (54), CRISPR-Cas systems rarely serve as an adaptive immune system protecting extant bacteria and archaea from infections with phage?

## Acknowledgements

This endeavor was supported by a grant from the Simons Foundation (396001), JG, and a grant from the US National Institutes of Health, R01-GM091875, BRL. This endeavor was supported by a grant from the Simons Foundation (396001), JG, and a grant from the US National Institutes of Health, R01-GM091875, BRL. We are grateful to Waqas Chaudhry for interesting and stimulating discussions and thank Ingrid McCall and Brandon Berryhill for help in preparing this manuscript and the editor and reviewers for useful comments and suggestions for improving the quality of the presentation.

## REFERENCES

1. Luria SE, Human ML. A nonhereditary, host-induced variation of bacterial viruses. J Bacteriol. 1952;64(4):557–69.

2. Arber W. Host-controlled modification of bacteriophage. Annual Review of Microbiology. 1965;19:365–78.

3. Barrangou R, Fremaux C, Deveau H, Richards M, Boyaval P, Moineau S, et al. CRISPR provides acquired resistance against viruses in prokaryotes. Science. 2007;315(5819):1709–12.

4. Wilson GG, Murray NE. Restriction and modification systems. Annu Rev Genet. 1991;25:585–627.

5. Pleska M, Lang M, Refardt D, Levin BR, Guet CC. Phage-host population dynamics promotes prophage acquisition in bacteria with innate immunity. Nat Ecol Evol. 2018;2(2):359–66.

6. Labrie SJ, Samson JE, Moineau S. Bacteriophage resistance mechanisms. Nat Rev Microbiol. 2010;8(5):317–27.

7. Hynes AP, Villion M, Moineau S. Adaptation in bacterial CRISPR-Cas immunity can be driven by defective phages. Nat Commun. 2014;5:4399.

8. Marraffini LA. CRISPR-Cas immunity in prokaryotes. Nature. 2015;526(7571):55–61.

9. Deveau H, Barrangou R, Garneau JE, Labonte J, Fremaux C, Boyaval P, et al. Phage response to CRISPR-encoded resistance in Streptococcus thermophilus. J Bacteriol. 2008;190(4):1390–400.

10. Childs LM, England WE, Young MJ, Weitz JS, Whitaker RJ. CRISPR-induced distributed immunity in microbial populations. PLoS One. 2014;9(7):e101710.

11. Levin BR, Moineau S, Bushman M, Barrangou R. The population and evolutionary dynamics of phage and bacteria with CRISPR-mediated immunity. PLoS Genet. 2013;9(3):e1003312.

12. Paez-Espino D, Morovic W, Sun CL, Thomas BC, Ueda K, Stahl B, et al. Strong bias in the bacterial CRISPR elements that confer immunity to phage. Nat Commun. 2013;4:1430.

13. Paez-Espino D, Sharon I, Morovic W, Stahl B, Thomas BC, Barrangou R, et al. CRISPR immunity drives rapid phage genome evolution in Streptococcus thermophilus. MBio. 2015;6(2).

14. Weissman J, Holmes R, Barrangou R, Moineau S, Fagan W, Levin B, et al. Immune loss as a driver of coexistence during host-phage coevolution. ISME. 2018:1–13.

15. van Houte S, Buckling A, Westra ER. Evolutionary Ecology of Prokaryotic Immune Mechanisms. Microbiol Mol Biol Rev. 2016;80(3):745–63.

16. Mayer A, Mora T, Rivoire O, Walczak AM. Diversity of immune strategies explained by adaptation to pathogen statistics. Proc Natl Acad Sci U S A. 2016;113(31):8630–5.

17. Chao L, Levin BR, Stewart FM. A complex community in a simple habitat: an experimental study with bacteria and phage. Ecology. 1977;58:369–78.

18. Wei Y, Kirby A, Levin BR. The population and evolutionary dynamics of Vibrio cholerae and its bacteriophage: conditions for maintaining phage-limited communities. The American Naturalist. 2011;178(6):715–25.

19. Wei Y, Ocampo P, Levin BR. An experimental study of the population and evolutionary dynamics of Vibrio cholerae O1 and the bacteriophage JSF4. Proc Biol Sci. 2010;277(1698):3247–54.

20. Bohannan BJ, Lenski RE. Linking genetic change to community evolution: insights from studies of bacteria and bacteriophage. Ecology letters. 2000;3(4):362– 77.

21. Koskella B, Parr N. The evolution of bacterial resistance against bacteriophages in the horse chestnut phyllosphere is general across both space and time. Phil Trans R Soc B. 2015;370(1675):20140297.

22. Lenski RE, Levin BR. Constraints on the coevolution of bacteria and virulent phage : a model, some experiments, and predictions for natural communities. American Naturalist. 1985;125:585–602.

23. Meyer JR, Dobias DT, Weitz JS, Barrick JE, Quick RT, Lenski RE. Repeatability and contingency in the evolution of a key innovation in phage lambda. Science. 2012;335(6067):428–32.

24. Levin BR, Stewart FM, Chao L. Resource - limited growth, competition, and predation: a model and experimental studies with bacteria and bacteriophage. American Naturalist. 1977;977:3–24.

25. Monod J. The growth of bacterial cultures. Annual Review of Microbiology. 1949;3:371–94.

26. Stewart FM, Levin BR. Resource partitioning and the outcome of interspecific competition: a model and some general considerations. American Naturalist. 1973;107:171–98.

27. Chaudhry WN, Pleska M, Shah NN, Weiss H, McCall IC, Meyer JR, et al. Leaky resistance and the conditions for the existence of lytic bacteriophage. PLoS Biol. 2018;16(8):e2005971.

28. Levin BR, Stewart FM, Chao L. Resource-limited growth, competition, and predation: a model and experimental studies with bacteria and bacteriophage. The American Naturalist. 1977;111(977):3–24.

29. Roach DR, Leung CY, Henry M, Morello E, Singh D, Di Santo JP, et al. Synergy between the Host Immune System and Bacteriophage Is Essential for Successful Phage Therapy against an Acute Respiratory Pathogen. Cell host & microbe. 2017;22(1):38-47.e4.

30. Schrag S, Mittler JE. Host parasite coexistence: the role of spatial refuges in stabilizing bacteria-phage interactions. American Naturalist. 1996;148:438–377.

31. Levin B. Restriction-modification and the maintenance of genetic diversity in bacterial populations. In: Nevo E, Karlin S, editors. Proceedings Conference on Evolutionary Processes and Theory. New York: Academic Press; 1986.

32. Levin BR. Frequency-dependent selection in bacterial populations. Philosophical Transactions of the Royal Society of London - Series B: Biological Sciences. 1988;319(1196):459–72.

33. Korona R, Levin BR. Phagemediated selection and the evolution and maintenance of restriction-modification. Evolution. 1993;47:556–75.

34. Levin BR. Nasty viruses, costly plasmids, population dynamics, and the conditions for establishing and maintaining CRISPR-mediated adaptive immunity in bacteria. PLoS Genet. 2010;6(10):e1001171.

35. Childs LM, Held NL, Young MJ, Whitaker RJ, Weitz JS. Multiscale model of CRISPR-induced coevolutionary dynamics: diversification at the interface of Lamarck and Darwin. Evolution. 2012;66(7):2015–29.

36. Brockhurst MA, Buckling A, Rainey PB. The effect of a bacteriophage on diversification of the opportunistic bacterial pathogen, Pseudomonas aeruginosa. Proceedings of the Royal Society of London B: Biological Sciences. 2005;272(1570):1385–91.

37. Koskella B. Phage-mediated selection on microbiota of a long-lived host. Current Biology. 2013;23(13):1256–60.

38. van Houte S, Ekroth AK, Broniewski JM, Chabas H, Ashby B, Bondy-Denomy J, et al. The diversity-generating benefits of a prokaryotic adaptive immune system. Nature. 2016;532(7599):385–8.

39. Weigle JJ, Delbruck M. Mutual exclusion between an infecting phage and a carried phage. J Bacteriol. 1951;62(3):301–18.

40. Pleska M, Qian L, Okura R, Bergmiller T, Wakamoto Y, Kussell E, et al. Bacterial Autoimmunity Due to a Restriction-Modification System. Curr Biol. 2016;26(3):404–9.

41. Qian L, Kussell E. Evolutionary dynamics of restriction site avoidance. Physical review letters. 2012;108(15):158105.

42. Kruger DH, Bickle TA. Bacteriophage survival: multiple mechanisms for avoiding the deoxyribonucleic acid restriction systems of their hosts. Microbiological Reviews. 1983;47(3):345–60.

43. Korona R, Korona B, Levin BR. Sensitivity of naturally occurring coliphages to type I and type II restriction and modification. Journal of General Microbiology. 1993;139(Pt 6):1283–90.

44. Pleska M, Guet CC. Effects of mutations in phage restriction sites during escape from restriction-modification. Biol Lett. 2017;13(12).

45. Kussell E, Leibler S. Phenotypic diversity, population growth, and information in fluctuating environments. Science. 2005;309(5743):2075–8.

46. Palmer KL, Gilmore MS. Multidrug-resistant enterococci lack CRISPR-cas. MBio. 2010;1(4).

47. Garrett RA, Shah SA, Vestergaard G, Deng L, Gudbergsdottir S, Kenchappa CS, et al. CRISPR-based immune systems of the Sulfolobales: complexity and diversity. Biochemical Society transactions. 2011;39(1):51–7.

48. Bradde S, Vucelja M, Tesileanu T, Balasubramanian V. Dynamics of adaptive immunity against phage in bacterial populations. PLoS computational biology. 2017;13(4):e1005486.

49. Campbell A. Conditions for the existence of bacteriophage. Evolution. 1961;15:153–65.

50. Lenski RE. Experimental studies of pleiotropy and epistasis in Escherichia coli. I. Variation in competitive fitness among mutants resistant to virus T4. Evolution. 1988;42(3):425–32.

51. Bohannan BJ, Kerr B, Jessup CM, Hughes JB, Sandvik G. Trade-offs and coexistence in microbial microcosms. Antonie Van Leeuwenhoek. 2002;81(1-4):107–15.

52. Bohannan BJ, Lenski RE. Effect of resource enrichment on a chemostat community of bacteria and bacteriophage. Ecology. 1997;78(8):2303–15.

53. Westra ER, van Houte S, Oyesiku-Blakemore S, Makin B, Broniewski JM, Best A, et al. Parasite exposure drives selective evolution of constitutive versus inducible defense. Current Biology. 2015;25(8):1043–9.

54. Datsenko KA, Pougach K, Tikhonov A, Wanner BL, Severinov K, Semenova E. Molecular memory of prior infections activates the CRISPR/Cas adaptive bacterial immunity system. Nat Commun. 2012;3:945.

